# Marker selection strategies for circulating tumor DNA guided by phylogenetic inference

**DOI:** 10.1101/2024.03.21.585352

**Authors:** Xuecong Fu, Zhicheng Luo, Yueqian Deng, William LaFramboise, David Bartlett, Russell Schwartz

## Abstract

**Motivation:** Blood-based profiling of tumor DNA (“liquid biopsy”) has offered great prospects for non-invasive early cancer diagnosis, treatment monitoring, and clinical guidance, but require further advances in computational methods to become a robust quantitative assay of tumor clonal evolution. We propose new methods to better characterize tumor clonal dynamics from circulating tumor DNA (ctDNA), through application to two specific questions: 1) How to apply longitudinal ctDNA data to refine phylogeny models of clonal evolution, and 2) how to quantify changes in clonal frequencies that may be indicative of treatment response or tumor progression. We pose these questions through a probabilistic framework for optimally identifying maximum likelihood markers and applying them to characterizing clonal evolution.

**Results:** We first estimate a distribution over plausible clonal lineage models, using bootstrap samples over pre-treatment tissue-based sequence data. We then refine these lineage models and the clonal frequencies they imply over successive longitudinal samples. We use the resulting framework for modeling and refining tree distributions to pose a set of optimization problems to select ctDNA markers to maximize measures of utility capturing ability to solve the two questions of reducing uncertain in phylogeny models or quantifying clonal frequencies given the models. We tested our methods on synthetic data and showed them to be effective at refining distributions of tree models and clonal frequencies so as to minimize measures of tree distance relative to the ground truth. Application of the tree refinement methods to real tumor data further demonstrated their effectiveness in refining a clonal lineage model and assessing its clonal frequencies. The work shows the power of computational methods to improve marker selection, clonal lineage reconstruction, and clonal dynamics profiling for more precise and quantitative assays of tumor progression.

**Availability:** https://github.com/CMUSchwartzLab/Mase-phi.git.

## Introduction

The discovery of circulating free DNA (cfDNA) in human blood and the observation that tumor-derived cfDNA may occur at greatly elevated levels compared to DNA of healthy cells — due to elevated release of tumor cell DNA, abnormal clearance for DNA debris from cell death, or circulating tumor cells in blood — established the potential for liquid biopsy, i.e., blood-based profiling of solid tumor genomics (Crowley et al., 2013; Wan et al., 2017). The prospect of rapid, non-invasive profiling of tumor states offers many possibilities for improving cancer diagnosis and treatment (Cescon et al., 2020) including early prognosis (Phallen et al., 2017b; Connal et al., 2023) and detecting residual disease and relapse(Mattox et al., 2019; Ignatiadis et al., 2021). Liquid biopsy methods have now been studied widely in various cancer types (Maia et al., 2020; Kemper et al., 2023).

The technology has technical limitations, however, mainly due to the challenge of separating tumor signals from the influence of much larger numbers of healthy cells and the consequent need for highly sensitive genomic assays. Deep sequencing on liquid biopsy samples with low signal-to-noise ratio is one option but can be too costly and time-consuming for repeated use. As a a result, alternative molecular testing methods have been used, including multiplex-PCR (Abbosh et al., 2017) and droplet digital PCR (ddPCR) (Huerta et al., 2021), as well as strategies for enriching for tumor DNA with targeted sequencing (Kurtz et al., 2021; Phallen et al., 2017a). These technologies offer a path to highly sensitive quantitation of somatic variants found at low levels in the blood, although with the tradeoff of allowing for profiling of relatively few pre-selected markers.

Despite its broad potential, current clinical application of liquid biopsy has primarily been for prognosis or recurrence detection (Reinert et al., 2019; Sanz-Garcia et al., 2022), rather than more precise quantitative analysis of tumor genetics. While there is now a rich literature on characterizing tumor evolutionary trajectories from numerous forms of genomic assays (c.f., (Beerenwinkel et al., 2015; Schwartz and Schäffer, 2017)), the need to work typically with low precision or relatively limited blood-based marker sets makes it infeasible to incorporate longitudinal blood samples in a straightforward way into current methods for multi-sample tumor phylogenetics. Yet there is also little work to date on developing new classes of inference method suitable for effectively bringing liquid biopsy into tumor phylogeny models. One notable exception has been recent work from the TRACERx Consortium using a tumour-specific phylogenetic method to profile ctDNA from non-small-cell lung cancer patients (Abbosh et al., 2017). That study inferred a base phylogenetic tree for each patient with primary multi-regional sequencing and then used PCR on liquid biopsy samples, preoperative and post-operative, to track clonal and subclonal populations. The same team later investigated a larger cohort with metastasis and used a new PCR technique and a bioinformatics tool tailored for ctDNA to track the clonal lineages longitudinally (Abbosh et al., 2023). Their work showed that liquid biopsy samples can identify mutations arising in distinct subclones and characterize clonal population changes in metastases or relapses, provided they can draw on an accurate model of clonal lineages. However, much remains unaddressed with regard to how to identify optimal markers for use in such analyses and how to use these most effectively to determine the clonal lineage model and how its population frequencies evolve over time.

The present work is aimed at developing methods to better characterize clonal dynamics of tumors from liquid biopsy data. We focus on challenges that have not, to our knowledge, been addressed in prior work. First, we examine the question of how we can leverage liquid biopsy data using small marker sets to refine phylogenetic models from the primary tumor so as to correct errors, reduce uncertainty in inference, or expand a tree to accommodate variants or clones not seen in earlier samples. Second, we consider the question of optimal marker selection: for typical scenarios in which one must select a small subset of markers to profile with high sensitivity, which markers are likely to be most informative? We examine this question for selecting markers to optimally refine the tumor phylogeny model and for measures of optimally characterizing changes in clonal frequency or tumor heterogeneity over time. We then show on simulated and real data that our methods allow one to apply liquid biopsy so as to accurately capture dynamics of clonal population changes in tumors, with potential application to various tasks in tumor diagnostics and clinical decision-making.

## Method

In this section, we consider variants of the problem of marker selection for liquid biopsy. For each, we assume we need to select a small marker set for high-precision assays, such as by ddPCR. Our goal is to develop personalized assays that allow rapid longitudinal corrections on a patient-specific basis. Note that we solve the problem for the general case of assuming that we might select from any observed marker for each specific subject, however the problem is conceptually the same if we are limited to choosing a subset of markers from a larger predefined set for which PCR probes are already available. We first consider the problem of choosing markers so as to refine a phylogenetic model and reduce uncertainty in clonal lineage inference. We then consider selection with the goal of characterizing changes in clonal frequencies given a known tree. The overall workflow for simultaneously addressing these questions is shown in Fig 1.

**Fig. 1:**
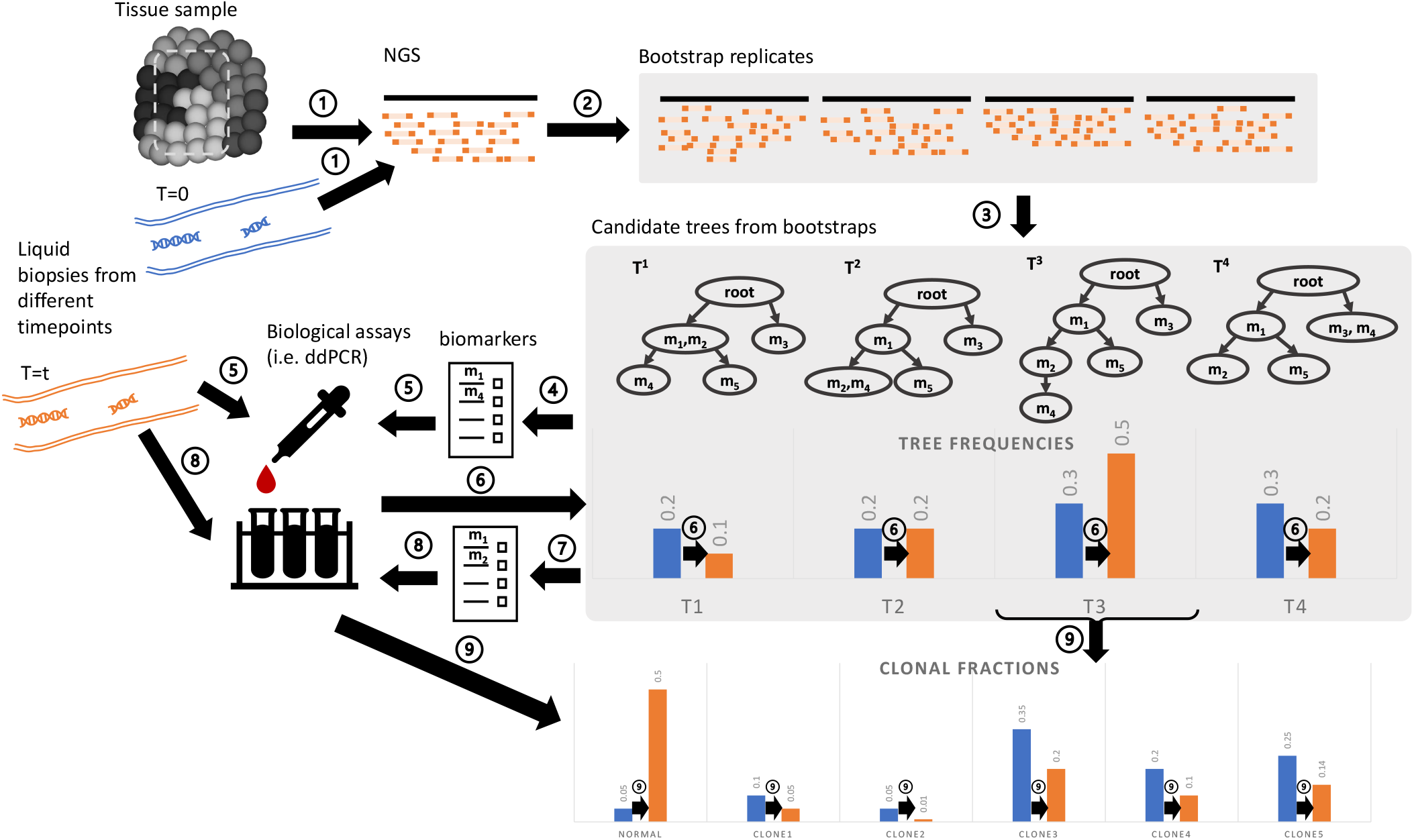
The overall inference pipeline. (1). We assume we have first sequenced tissue and liquid biopsy sample(s), obtaining reference germline and ctDNA. (2) We create bootstrapped samples over reads for each sequence set. (3) We infer a set of possible trees from the bootstrapped samples, serving as an estimated empirical tree distribution. (4) We then seek a set of optimal biomarkers of mutations to best reduce the tree uncertainty and (5) apply these in biological assays (e.g. ddPCR). (6) We then use the results of these assays to update the empirical tree distributions. (7) We further seek a set of optimal biomarkers to track subclone frequencies efficiently and (8) assay these biomarkers. (9) Finally, we then use the results of the assays to estimate clonal fractions at each sampled timepoint.

### Selecting mutation markers to minimize uncertainty in phylogenetic inference

We first consider the problem of choosing liquid biopsy markers so as to reduce uncertainty in the tree inference. Due to the low signal-to-noise of liquid biopsy samples, we can expect genomic measurements from liquid biopsy to yield poor results with standard phylogenetic inference tools. Fig 2 demonstrates this with simulated data. As a result, we assume there will be high uncertainty in tree inference and pose phylogenetic inference in terms of distributions of trees rather than a single optimum. For the present purpose, we estimate this distribution through bootstrapping over sequence reads, an approach chosen because it allows us to use existing tumor phylogeny methods that are designed to return a single optimal tree. Bayesian phylogeny methods might provide a more principled alternative than our bootstrapping approach to capture the initial tree distribution, although designing an efficient Bayesian sampler for non-trivial tumor phylogeny models is a challenging problem in itself. Once we have an initial tree density, we then seek in part to select liquid biopsy markers that will allow us to reduce uncertainty in the tree inference by facilitating comparisons that can reject some subset of the topologies. For this purpose, we want to find marker sets that are to best distinguish between possible high-frequency models in the initial tree distribution.

**Fig. 2:**
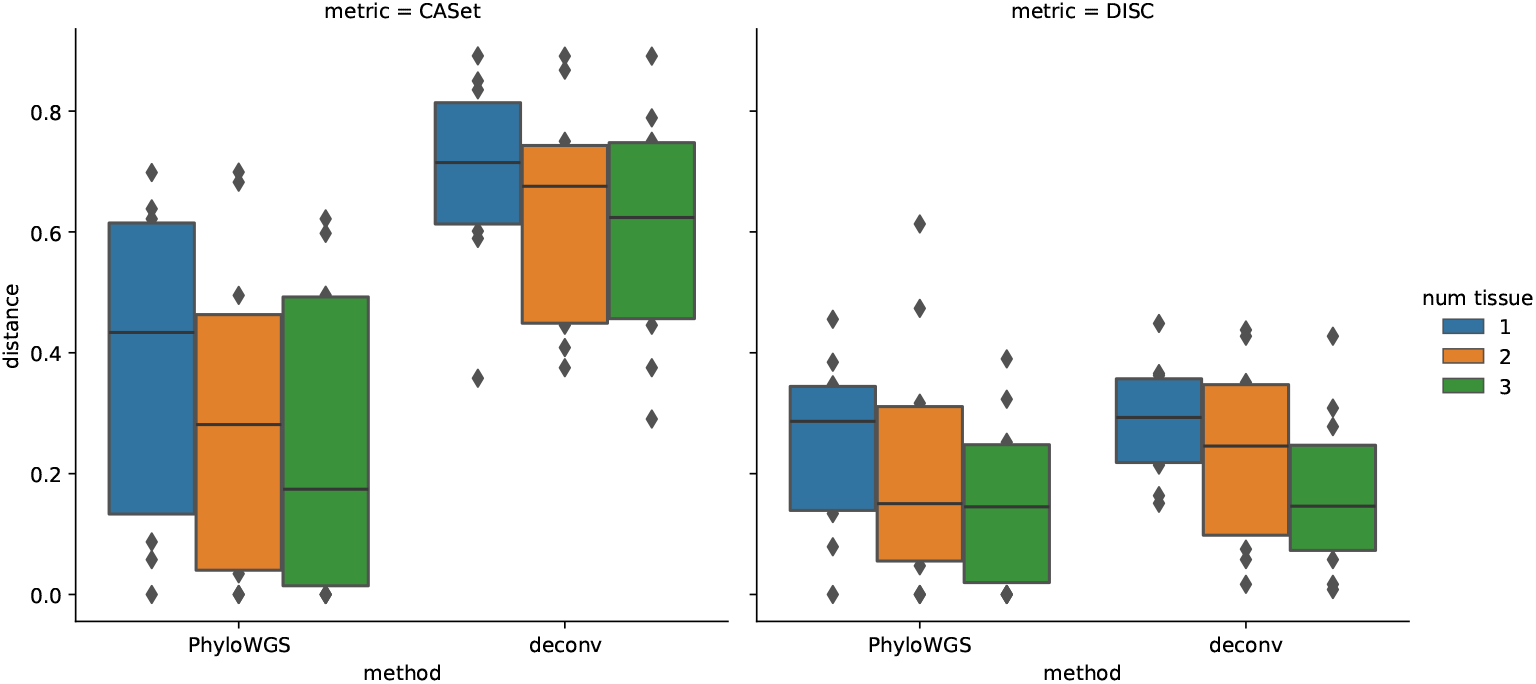
Assessment of accuracy of tree inference using paired liquid biopsy and multi-regional tissue samples. The figure shows distance between true and inferred trees as function of number of samples for the PhyloWGS and deconv methods. (a) CASet distance (Dinardo et al., 2020). (b) DISC distance (Dinardo et al., 2020).

### Problem formulation

We first establish a general probabilitic framework for defining a tree density and posing optimal marker selection problems over it. In subsequent sections, we adapt this to distinct solutions to the problem. At a high level, each variant of the method described below works by estimating a density over trees and optimizing marker selection for a desired objective over that density. The notation below is a formalization of this basic idea, then adapted to different problem assumptions and objectives.

Given a candidate (bootstrapped) tree set 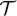 = {*T*^*k*^}_*k*=1,…,*K*_ we define a clonal tree structure matrix *E*^*k*^ that specifies possible structures for a set of defined tumor clones. For each tree *k*, 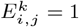 if clone (tree node) *i* is the parent of node *j* and otherwise 0. We define a mutation assignment matrix *M*^*k*^ where 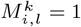 if mutation *g*_*l*_ belongs to node *i* for each tree *k* and otherwise 0. We define a clonal frequency array 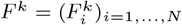, encoding the variant allele frequency (VAF) of each of *N* observed mutations in each clone *i* for each tree *k*. We further define the set of mutations 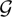 = {*g*_*l*_}_*l*=1…*N*_. We want to select *n ≤ N* gene markers 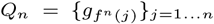 by finding a mapping *f*_*n*_such that the 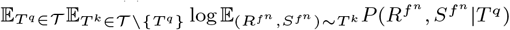 Is minimized, where 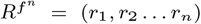 is a family of random variables describing ddPCR read counts for a set of probes and 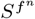 is a binary matrix mapping biomarkers to clones in the tree structure for an underlying ground truth tree *T*^*k*^. In the model, we assume that the generation of read counts is i.i.d. and independent from the generation of the tree structures for simplicity of calculation.

We split the likelihood of a given set of data into the probability of observing the read counts and the probability of observing the tree structures. Assume that the read count for each gene marker 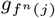 is *r*_*j*_. Then letting *M*^*k*^(*l*) = *i* when 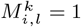:

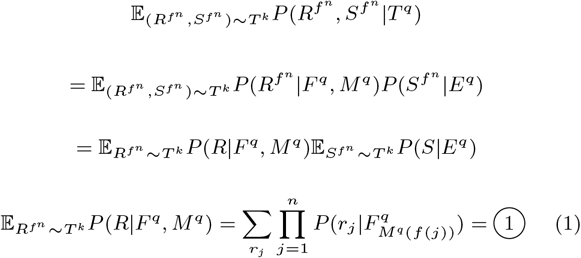

Since the ddPCR is assumed to have high sequencing depth, we assume that the probability of observing the read counts is close to a normal distribution instead of a binomial for computational convenience. Assume the read depth is *D*. Then assuming 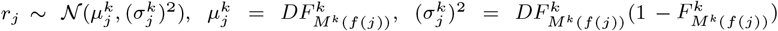

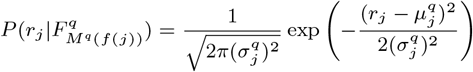

where 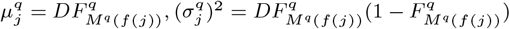

Then:

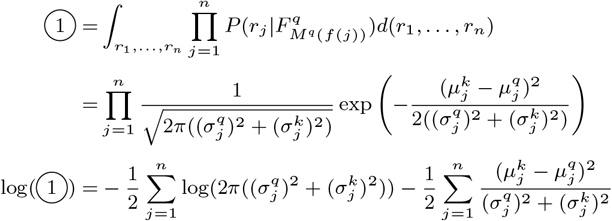

For the probabilistic modeling for structural perturbations, we use the ancestor-descendant distance (Govek et al., 2020) to model *Q* the distance between subtrees. We define a subtree 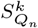 of *T*^*k*^ given a set of marker genes *Q*_*n*_ using their ancestor-descendent matrix *A*. For *T*^*k*^, 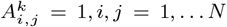 if mutation *i* is the ancestor of *j*. Then for 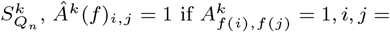 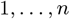. For each unit change of AD distance, the probability is *λ*. Therefore

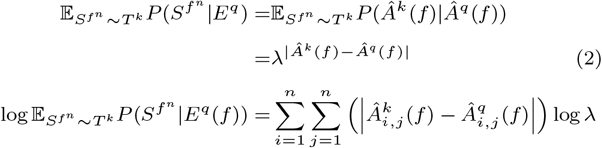

After combining the two components, we get an objective function that we seek to minimize in marker selection:

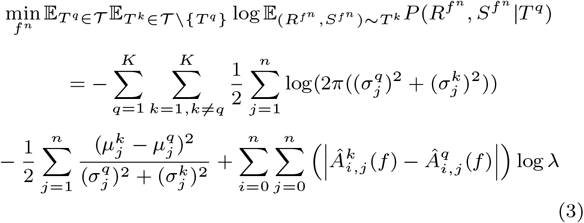

Effectively, this objective function provides a way of evaluating the utility of a given marker set for reducing a measure of uncertainty in expectation over the tree density and sampling of sequence reads.

### Refining tumor phylogenetic tree distribution after liquid biopsy assays on selected markers

For the ddPCR counts tested for selected marker genes *g*_1_ and *g*_2_, suppose without loss of generality that *g*_1_ is the ancestor of *g*_2_ in a proposed tree structure. We then refine the tree structure by testing which trees are consistent with all the possible relationships among *g*_1_ and *g*_2_ consistent with the new data. While one might build this into the tree likelihood and solve *de novo* for the tree density, for efficiency reasons we instead apply new marker data by posing it as a problem of refining an existing density over trees. We consider two variants: first posing this as a simpler statistical hypothesis testing problem and then as a series of Bayesian updates.

We assume in the discussion below that the read depths for each tree are described by binomial variables based on the allele frequencies and overall read depth: 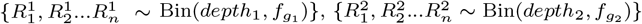.

### Significance testing method

One straightforward approach to using ctDNA to refine a tree distribution is to accept or reject potential candidate trees based on whether the measured marker frequencies are plausibly consistent with a given tree topology. We assume here that frequencies of clones may change over time in ways that reveal some trees to be implausible that were initially plausible, but the set of clones and their tree topology are unchanged over the course of the follow-up. Given a pair of markers, we can pose the question of whether their read counts at a specific point in time are consistent with a given tree topology as a statistical hypothesis test, here using a Wald test with the test statistic

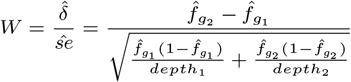

with size *α*. We reject the hypothesis when *W > z*_*α*_, with *α* = 0.05 in the present work. For each pair of markers, we perform the Wald test with a Bonferroni-corrected *α* and then remove from the density all tree structures that are rejected by any pairwise marker test. Repeating this for all pairs of markers in a set then gives a general test to reject a portion of the tree density and lead to a refined density consistent with both the original sequence and subsequent ctDNA data. This test may thus be applied serially for multiple longitudinal assays, provided the original density was sufficiently well sampled that the correct tree is found within it.

### Bayesian update method

Statistical hypothesis testing might be too strict for some uses, in allowing us only to accept or reject a given tree and the latter only with compelling evidence. ctDNA data might still give evidence for or against certain trees without being able to definitively accept or reject them. We therefore also develop a Bayesian approach to capture more nuanced changes in our inferred tree distribution by updating the weights for each possible topology in the distribution to reflect its plausibility given all of the data seen to date. Since we used bootstrapping trees to approximate the tree distribution from the observed data, we define the initial weight of each tree structure to be the count of that tree structure observed in the bootstrapping. Note that these counts are normalized to produce a probability density over trees but are represented initially as integers here. We keep our prior notation *S* to represent the tree structure, *R*^0^ for the observed read counts from the original primary tissue sequencing, and *R*^1^ for the observed PCR counts from the liquid biopsy samples. Then:

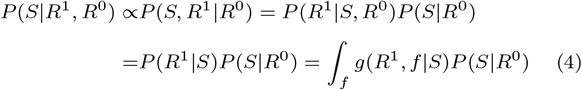

where *P*(*S*|*R*^1^, *R*^0^) defines the updated weights, *P*(*S*|*R*^0^) is the original weights,*f* describes the possible VAFs for the mutation markers from liquid biopsy, and *g*(*R*^1^, *f* |*S*) is the probability density of a particular set of clonal VAFs *f* and the corresponding read counts, given a structure *S*.

We develop the special case of two markers to derive the basic method for refining the tree topology and for use in subsequent illustration. We can generalize the method to multiple independent marker pairs through sequential application of pairwise updates. A more rigorous but tractable generalization to *k* markers for arbitrary *k* is less trivial and left as an exercise for future work.

For two markers, there are four types of relationship those markers might take on in a tree structure: (a) marker 1 is an ancestor of marker 2, which means that *f*_1_ *> f*_2_. (b) marker 2 is an ancestor of marker 1, meaning *f*_1_ < *f*_2_, (c) marker 1 and marker 2 belongs to the same clone, meaning *f*_1_ = *f*_2_ = *f*, (d) marker 1 and marker 2 belong to different branches of the tree, meaning *f*_1_ + *f*_2_ < 1. Therefore, let 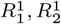 be the read counts of marker 1 and 2 from liquid biopsy and 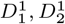 be the read depths. Then:

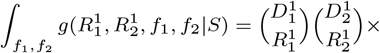

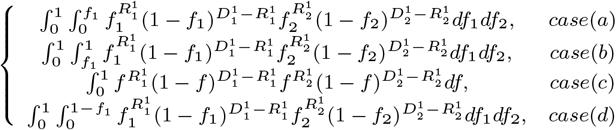

Equation 4 provides a formula to update the weight of each possible tree by multiplying each by the above integral. We then normalize the overall updated weights of all possible structures to sum to 1 in order to yield an updated tree density. We perform marker selection for the purpose of optimal tree refinement by integer linear programming (ILP) over possible choices of markers and over trees in the tree density to find a marker set optimizing for the objective function (3), as described in more detail below.

### Selecting mutation markers to track the changes in subclonal populations

We next consider another important criterion for marker selection: effectively tracking changes in subclonal populations. We would typically assume that our purpose in conducting liquid biopsy is to track changes in clonal frequencies or changes in overall heterogeneity that might be indicative, e.g., of recurrence after treatment, growth of a resistant clone, or metastasis. Intuitively, to optimize for this criterion, we want to avoid choosing redundant markers but rather find a set of markers that are distributed across the phylogeny so as to provide as much power as possible to monitor changes in distinct clonal frequencies.

To facilitate our explanation, we first derive a method for solving this problem under the assumption that we have determined a specific lineage tree and we want to select a set of markers to accurately track the clonal dynamics of the tumor as a whole. We later generalize that to the actual case where we assume a distribution over trees rather than a single known tree.

### Deterministic Trees

#### Problem Statement

Given the same input as 2.1, and an index 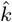 which indicates the most likely tree, select *n* gene markers {*g*_*f*(*j*)_}_*j*=1…*n*_ by finding a mapping *f*(*j*) such that the sum of weighted tracked clones by the markers in 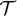 is maximized.

Here, we set the weights to be the estimated clonal fractions from the previous time point, posing the problem so as to maximize the estimated fraction of the tumor tracked. However, the weights here could be any arbitrary design, for example if we wanted to bias the selection to favor particular probes based on measures of their expected clinical utility, ease of probe design, preference for probes already available, or some other application of expert knowledge.

For computational convenience, we define the weight for a tree node to be the weight of each the mutations first appearing in that node. We then create a clonal frequency array 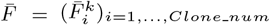 from the most recent estimate of the variant allele fractions for each clone 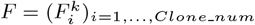, where 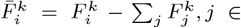 Children(*i*). In normal use in longitudinal sampling, these weights would then update with each longitudinal time point to provide a best guess as to the weights at the next time point. *S*^*k*^ is the matrix of pairwise “same-node” relationship of the *k*^*th*^ bootstrapped tree. 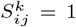 if mutations *i* and *j* belongs to the same node in the *k*^*th*^ tree and 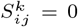 otherwise, where *i > j*.

We further define a binarization operation 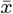 as follows

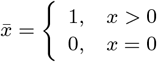

We then define a binary output vector *z* to identify the chosen markers as above. Let 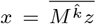, *x* be an array where *x*_*j*_ = 1 indicates the node *j* has been tracked and *x*_*j*_ = 0 if not. 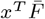 is the total proportion of the tracked clones that we minimize.

Measuring clonal frequencies is not entirely straightforward, though, because the mutations acquired in any node will be inherited by its descendent nodes. Therefore we can only identify the actual frequency of a given clone by measuring a marker of that clone as well as markers of its children’s clones. We call this the “complete information assumption” and call the previously illustrated scenario the “partial information assumption”. Under the complete information assumption, we create a matrix 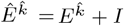 mapping clones to mutations whose VAFs would allow us to identify the clonal fraction. We define a pairwise product ⊙ between matrix *A* and array *b* such that (*A* ⊙ *b*)_*ij*_ = *a*_*ij*_ ∗ *b*_*j*_. We define a row-wise sum of a matrix *A* as *σ*(*A*) where *σ*(*A*)_*i*_ = ∑_*j*_ *a*_*ij*_. We can then pose the problem of finding the optimal marker set to correspond to solving the constrained optimization problem max_*z*_ 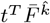 such that 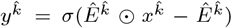 where 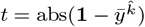.

### Candidate tree set

We next extend the simplified model, which assumes a known tree, to consider uncertainty in tree inferences, in which we assume we have a density over either a subset of candidate trees or the full tree space. Assume that 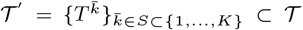 is the tree set over which we want to optimize. Our objective function would be max_*z*_ 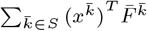, where 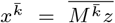 under the partial information assumption. Under the complete information assumption, the constrained optimization problem is transformed to max_*z*_ 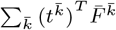 such that 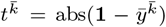 where 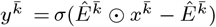 similar to the deterministic tree case.

### Tracking tumor subclonal population using the selected markers

After refining the tumor phylogenetic tree distribution, we track the subclonal population by using the chosen markers. We infer the frequency of a clone using the mean VAF of markers appearing in the given clone minus the sum of mean VAFs of markers inferred for its child clones. We note that this model does assume that we are only choosing from markers in copy number neutral regions and thus can treat VAF as a proxy for cancer cell fraction (CCF).

### Simulations

We create simulated phylogenetic tree structures parameterized by the number of subclones and the maximum degree of each subclone to control how many child subclones a parent node can have, randomizing subclone distributions to set up the total tree structure. Then we use a beta distribution to generate true allele frequencies for each subclone in this tumor tree. Based on the assumption that clones will have different frequencies at different tumor sites or in tissue versus blood, we used a Dirichlet probability distribution to randomize clonal frequencies with tumor tissues. Yo mimic observations on real tumor and blood samples, we added a masking step so that only part of the total subclones that are nearer the root are observed in tissue samples and the rest can only be observed in liquid biopsy samples. We normalize the fractions of observed clones in each tissue sample so that the fractions add up to one. However, in liquid biopsy tumor blood samples, normal cells may dilute the mutated alleles in the blood, which will lead to a higher frequency in the normal cell. Therefore, we add a frequency of 0.9 to the normal cell with a subsequent Dirichlet random variable to randomize the rest of the frequency for each subclone. We use an additional Poisson random variable to randomly assign a number of mutations to each subclonal node in the tree.

Since we already simulate the total structure of the tree, the assignment of mutations, and their corresponding variant allele frequency, we are able to assign a depth of total reads for both tissue tumor samples and blood tumor samples. We then use a Poisson random variable to select for a total read count for each variant in each sample based on those numbers and use a binomial distribution with the probability equal to their variant allele frequency to determine the number of variant reads and reference reads.

To model sampled ddPCR measurements from this clonal growth model, we assign a number of droplets collected and determine the number of these detected to be the mutant based on the normalized real data, using the mean of droplets in the given real sample and the known variant allele frequency to simulate the number of droplets detected as mutant positive.

### Implementation of the optimization methods

All the probe optimization problems are solved by formulating them as integer linear programs (ILPs). We create a variable *z ∈* {0, 1}^*N×*1^ so that *z*_*i*_ = 1 if *i*th mutation is selected as a marker and *z*_*i*_ = 0 otherwise. We constrain 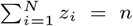 to impose the marker number constraint. We then optimize for each objective function with *z* as the variable, establishing an ILP. We use Python with Gurobi for the solver. We tested our problem with two tumor phylogeny methods, the MCMC-based method PhyloWGS (Deshwar et al., 2015) and a simplified version of our own ILP method, TUSV-ext (Fu et al., 2022), restricted to only consider SNVs. ILPs were all solved using Gurobi Ver.9 in the present testing.

## Results

### Simulated data

Before directly assessing our methods for marker selection and application, we first assessed how accurately we can determine phylogenies by different tumor phylogeny methods using simulated tissue and blood samples. Many tools have been developed for tumor phylogentics from tissue samples, including multiregional samples, yet none to our knowledge are designed specifically for including high noise-to-signal ratio liquid biopsy samples or have been tested on comparable data. We compared results using PhyloWGS (Deshwar et al., 2015) and an SNV-only version of TUSV-ext (Fu et al., 2022), which we refer to as `deconv`. We also applied two methods for measuring phylogenetic distance specialized to tumor phylogenies, CASet and DISC (Dinardo et al., 2020), for tree distance evaluation. For the present purposes, we assumed an initial liquid biopsy sample was sequenced along with one or more tissue samples. The results show that PhyloWGS yields good results even with only a single tissue sample, although accuracy is better the more tissue samples are included (Fig. 2). deconv performs poorly by the CASet measure, although comparably to PhyloWGS by the DISC measure. These results show that there is substantial room for improvement in tree inference, supporting the value of refining the tree model using subsequent ctDNA samples.

We then tested the whole analysis pipeline of Fig. 1 on the simulated data. We first generated 100 bootstrapped samples for each simulation case and inferred a tree for each bootstrap replicate. Since PhyloWGS performed better in the earlier tests, we use it subsequently to generate bootstrap trees. We then apply our marker selection methods on the bootstrap tree sets before applying the chosen markers to adjust the tree distribution according to the simulated ddPCR data. Our simulation results show that using even limited numbers of markers can shift the tree distribution towards the ground truth tree; almost all cases yield refined trees below the diagonal in 3(a) and (b), meaning that the updated weighted distance of the tree distribution to the ground truth tree is reduced by the update. The weights of the bests tree relative to alternatives are also improved, as we see in Fig 3(c). We compare each of these measures between markers chosen for the purpose of reducing uncertainty against those chosen for optimally characterizing clonal frequencies and against randomly selected markers. We see that markers chosen to optimize their ability to reduce uncertainty in the tree structure indeed perform substantially better at this task. Selecting markers for their utility in estimating clonal frequencies yields a marker set that performs more poorly at refining trees than one selected for the purpose of refining trees, as we would expect. Randomly chosen markers similarly perform poorly at updating tree topology, with instances where random markers outperform markers chosen for inferring clonal frequencies at the task of refining trees and vice versa.

**Fig. 3:**
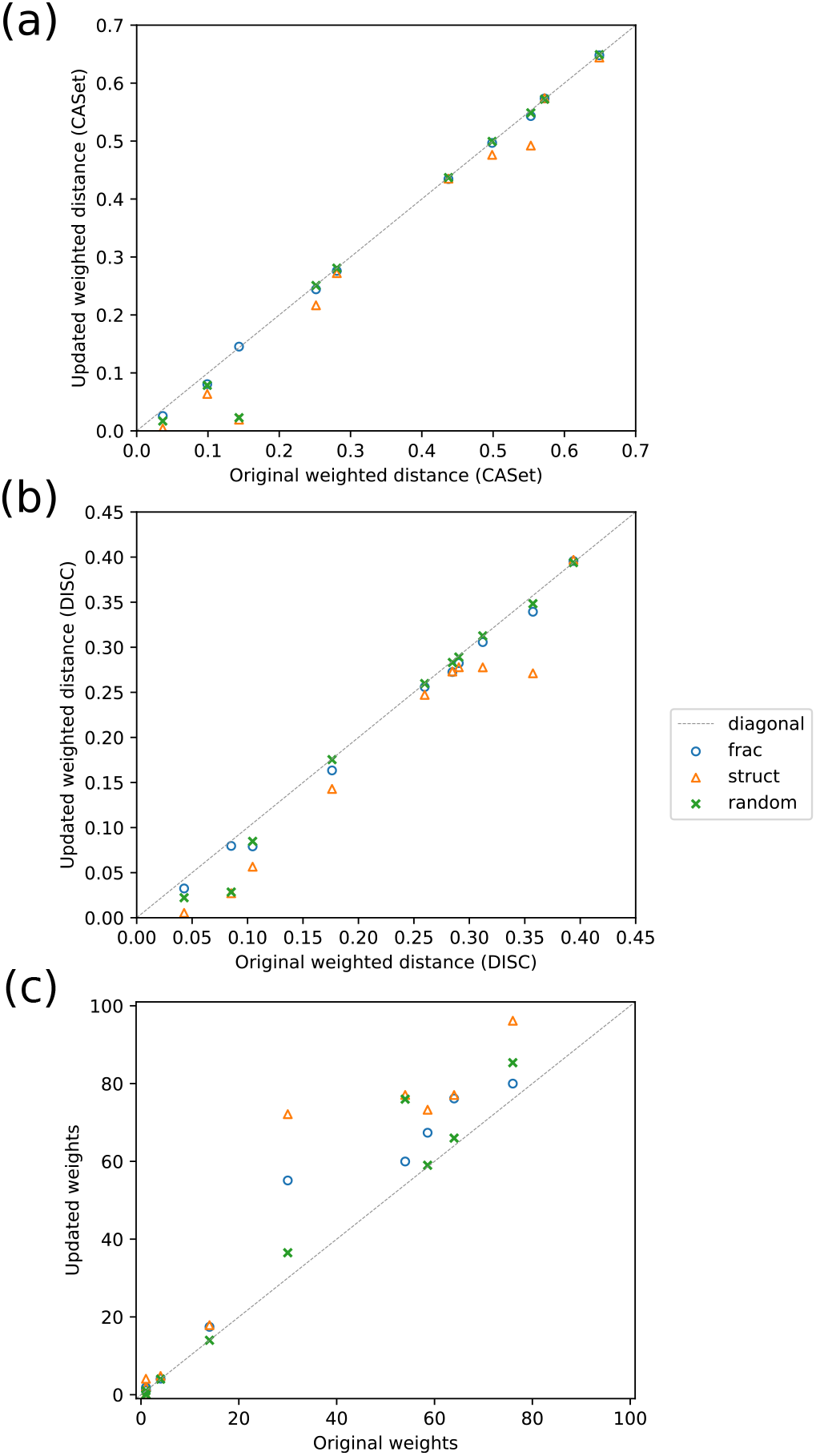
Simulation result from 10 random tumor clonal trees with 7 clones, mutation rate of 50 and mask proportion of 0.5 for tissue, with one liquid biopsy sample and two tissue samples. (a) The updated weighted distance after the tree adjustment versus the original weighted distance between the estimated tree and the ground truth tree, whose weights are the frequencies of each tree structure in the bootstrapped trees. CASet(Dinardo et al., 2020) is used as the distance metric. (b) The updated weighted distance after the tree adjustment versus the original weighted distance using DISC (Dinardo et al., 2020) as metric. (c) The updated weights versus the original weights for the best tree structure, with the lowest distance compared to the ground truth tree. We compared the three marker selection strategies: optimizing for inferring clonal fractions (frac), inferring tree structures (struct), and random selection (random).

We also evaluated the ability of the methods to monitor clonal fractions. We visualize the results for ten simulation cases in Fig 4, showing increasing ability to characterize clonal frequencies with larger numbers of optimally chosen markers. By comparison, inferences from randomly selected marker sets are typically unstable and require more markers to achieve minimal accuracy.

**Fig. 4:**
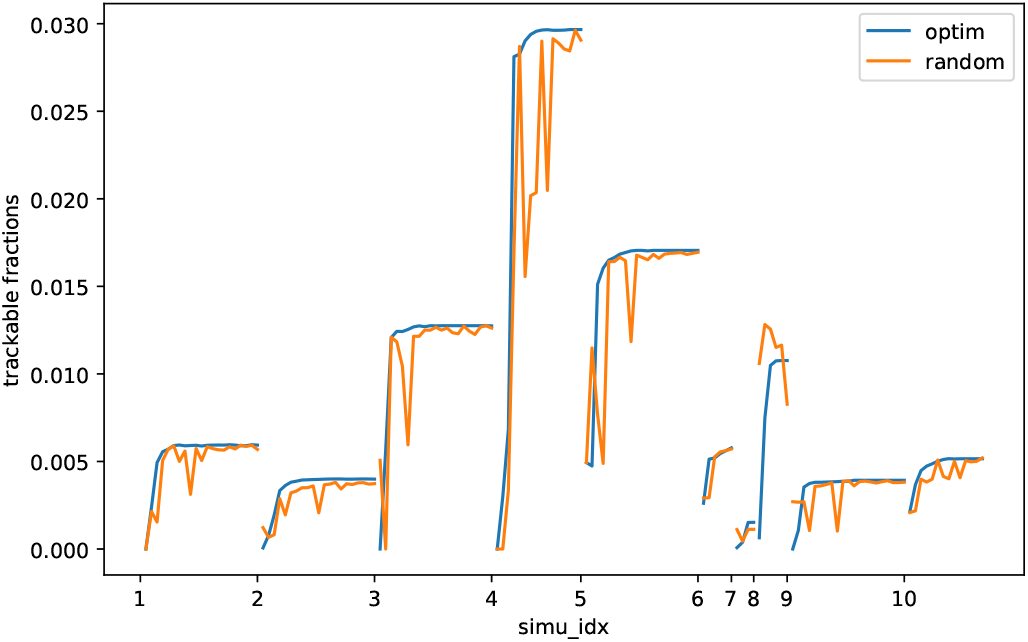
The trackable fractions of simulated tumor clonal population in noisy liquid biopsy samples. The lines show the change of trackable fractions, defined as the fraction of the total clonal frequency of clones that can be quantified with the optional marker set, as the selected marker number increases. We compare marker selection optimized for the task of maximizing trackable fraction against random marker selection.

### Application to a lung cancer case from the TRACERx study

Abbosh et al. (2017) assayed ctDNA for the first 100 TRACERx research participants, which Abbosh et al. (2023) expanded to additional participants. To demonstrate our methods, we selected one TRACERx case, CRUK0044, for which three primary tumor multi-regional samples were sequenced followed by six consecutive temporal liquid biopsy samples. Our simulation study suggested optimizing for tree structures provides a more informative basis for selecting markers than does optimizing for clonal frequencies, so we applied only tree structure based selection for this real data case. We iteratively selected markers for each time point, selecting a subset of a larger marker set actually profiled by the TRACERx study, and used these to update the empirical tree distributions (Fig. 5(a)). The blood samples (shown in the thick blue line) reinforced the the tree structure originally most frequently observed in bootstrap replicates from primary sequence, suggesting that the initial multiregional sequencing had likely given highest weight to the correct inference. Nonetheless, the ctDNA allowed us to increase our confidence in that inference by providing evidence against other possible topologies that were consistent with the original data. The first sample after the surgery tends to have the least ctDNA remaining, as we might expect, making the tree distribution adjustment less stable and temporarily leading to an inference that a lower-weight tree (green line in Fig 5(a)) was the most plausible. However, the tree distribution stabilizes after the first few blood draws with enhanced confidence in the initial most likely tree.

**Fig. 5:**
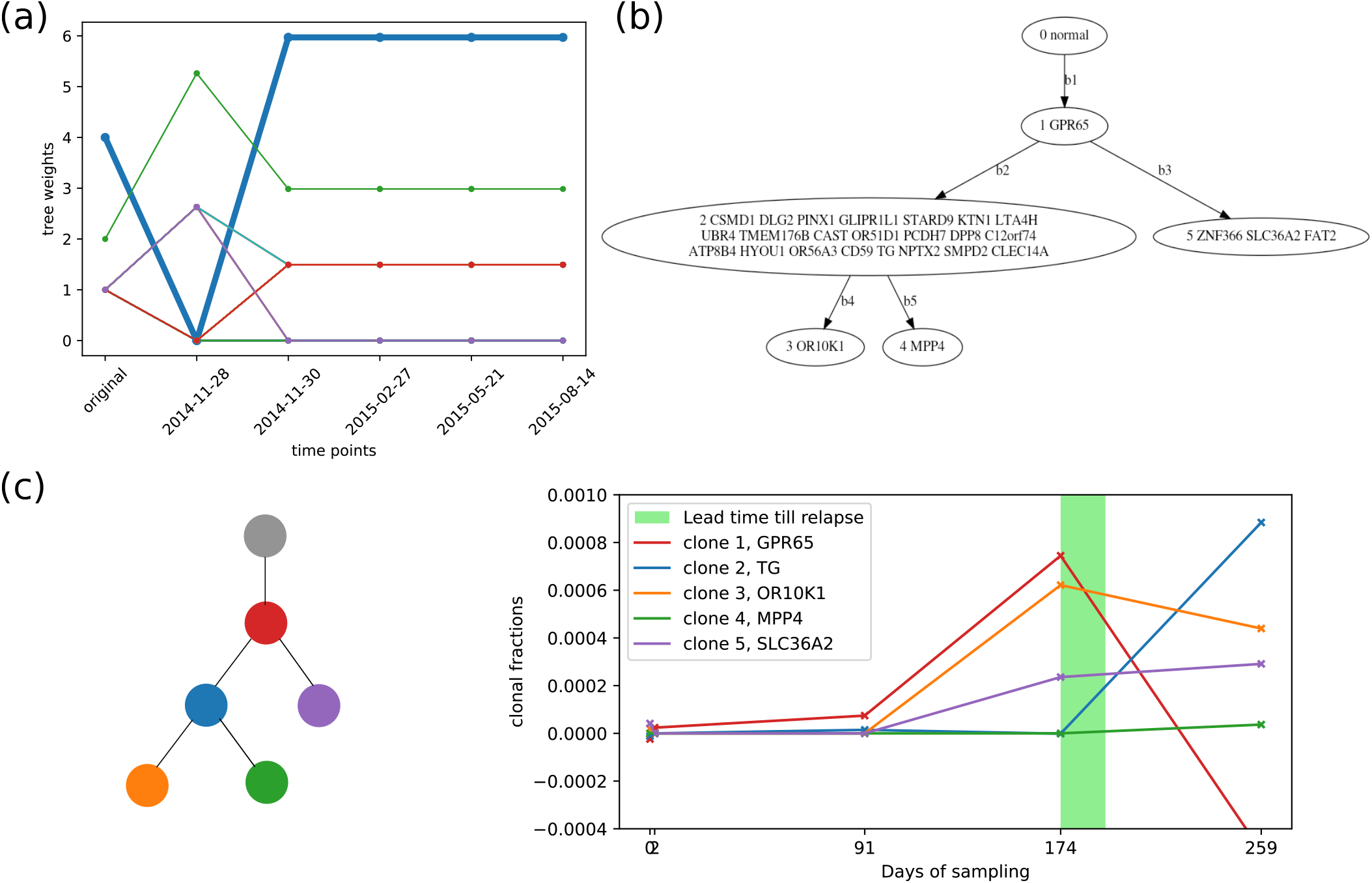
Results of applying our tree refinement and clonal tracking methods on the CRUK0044 sample from the TRACERx data. (a) Changes in tree weights for each topology identified in bootstrap sampling, after adjusting the tree distribution using the selected markers at each time point. (b) The inferred most likely tree after all serial samples, corresponding to the blue line in (a). (c) Inferred clonal frequencies as of each longitudinal sample derived from the selected marker set as of each day of sampling, with lines representing the clones color-coded as in the tree at left.

We then apply the second marker selection strategy to track the clonal subpopulations, using the most probable tree structure (Fig 5(b)) as the presumed correct tree. Since we have three multi-regional samples, we have an initial candidate list of markers from all three fraction sets. We select the top five markers among those occurring in at least two of the three sets. We tracked the fraction of each subclonal population for each time point of the collected liquid biopsy samples (Fig 5(c)). Our method is able to use just a small subset of markers to capture an overall trend of gradual and then accelerating expansion of several subclones after surgery. This trend was apparent in the sampling by day 91, although several of the clones that came to dominate in relapse were seemingly present at negligible levels in the blood until day 174. We also note a sharp rise of the initially rare subclone 5, which then becomes the dominant clone by day 259, post relapse. The data do reveal some imprecision in inferences, however, most prominently in the inference of a negative clonal frequency for the once-common clone 1. A negative clonal frequency indicates that markers identifying the descendants of clone 1 are observed at higher frequency than the markers for clone 1 itself, an impossible result if the tree and marker frequencies are exact (El-Kebir et al., 2015) but one that can occur due to imprecision in marker frequencies or error in the tree topology.

## Discussion

In this paper, we develop and apply a computational framework for interpreting ctDNA data for refining phylogenetic tree models and tracing clonal frequencies in tumor phylogenetic models. We further use this framework to develop optimization methods for selecting limited numbers of high-sensitivity markers for liquid biopsy in tracking cancer progression. We pose and solve for two optimization variants of the marker selection problem: selecting markers to refine tumor phylogeny models and to track clonal frequencies. Application to simulated data shows the methods yield good accuracy at both tasks with limited numbers of markers, substantially improved over markers selected randomly or for a different task. Application to real data further demonstrates the potential of the methods to bring liquid biopsy more effectively to studies of clonal lineage trees and to clinical applications that depend on precisely and quantitatively tracking changes in clonal dynamics over time.

The present work is largely a proof of concept of a general approach to phylogeny-assisted marker selection for liquid biopsy that might be extended in a number of ways. The modeling framework could be adapted to more sophisticated Bayesian models of tree space, for example to develop more principled but tractable strategies for handling larger marker sets. Other objective functions for defining optimal marker sets or interpreting their results might also be considered, more specifically tuned to specific clinical questions of interest. We also note that the present application focused on SNVs, but in future work it will be important to consider both structural variations (SVs), which are likely to be high impact and also make effective probes for tumor tracking, and copy number alterations (CNAs), which are often the mechanism of action of tumor driver genes and can confound interpretation of SNV VAFs. Finally, considerable work remains to be done to take advantage of the new capabilities informatics can enable for liquid biopsy to lead to improvements in the practice of public health interventions and clinical treatment of cancers.

## Acknowledgements

We thank Oana Carja and Thomas Rachman for helpful discussions.

